# Balancing selection of the Intracellular Pathogen Response in natural *Caenorhabditis elegans* populations

**DOI:** 10.1101/579151

**Authors:** Lisa van Sluijs, Kobus J. Bosman, Frederik Pankok, Tatiana Blokhina, Jop I. H. A. Wilten, Joost A. G. Riksen, Basten L. Snoek, Gorben P. Pijlman, Jan E. Kammenga, Mark G. Sterken

## Abstract

Genetic variation in host populations may lead to differential viral susceptibilities. Here, we investigate the role of natural genetic variation in the Intracellular Pathogen Response (IPR), an important antiviral pathway in the model organism *Caenorhabditis elegans* against Orsay virus (OrV). The IPR involves transcriptional activity of 80 genes including the *pals-*genes. We examine the genetic variation in the *pals*-family for traces of selection and explore the molecular and phenotypic effects of having distinct *pals*-gene alleles. Genetic analysis of 330 global *C. elegans* strains reveals that genetic diversity within the IPR-related *pals*-genes can be categorized in a few haplotypes worldwide. Importantly, two key IPR regulators, *pals-22* and *pals-25*, are in a genomic region carrying signatures of balancing selection, suggesting that different evolutionary strategies exist in IPR regulation. We infected eleven *C. elegans* strains that represent three distinct *pals-22 pals-25* haplotypes with Orsay virus to determine their susceptibility. For two of these strains, N2 and CB4856, the transcriptional response to infection was also measured. The results indicate that *pals-22 pals-25* haplotype shapes the defense against OrV and host genetic variation can result in constitutive activation of IPR genes. Our work presents evidence for balancing genetic selection of immunity genes in *C. elegans* and provides a novel perspective on the functional diversity that can develop within a main antiviral response in natural host populations.

## Introduction

Viral infections occur in natural populations of all organisms. Genetic variation can change host-virus interactions by altering coding sequences of protein products. Moreover, host-virus interactions can also be influenced by genetic variation due to altered gene copy numbers. Structural and regulatory genetic variation may both affect the viral susceptibility after infection, making some individuals within the population more resistant than others (Franco et al., 2013; van Sluijs et al., 2017; Piasecka et al., 2018; Wang et al., 2018). Presence of viruses can thereby select for beneficial genetic variants to remain present in the population (Enard et al., 2016; Wilke and Sawyer, 2016).

The nematode *Caenorhabditis elegans* and its natural pathogen Orsay virus (OrV) are used as a powerful genetic model system to study host-virus interactions (Félix et al., 2011). OrV is a positive-sense single-stranded RNA virus infecting *C. elegans* intestinal cells where it causes local disruptions of the cellular structures (Félix et al., 2011; Franz et al., 2013). This can result in lower fecundity in highly susceptible animals, but the infection does not affect lifespan (Félix et al., 2011; Sarkies et al., 2013). Three antiviral responses are known, of which RNA interference (RNAi) and uridylation both target viral RNA for degradation (Félix et al., 2011; Ashe et al., 2013; Sterken et al., 2014; Le Pen et al., 2018). The third response, the so-called Intracellular Pathogen Response (IPR), is thought to relieve proteotoxic stress from infection by OrV and other intracellular pathogens (Bakowski et al., 2014; Reddy et al., 2017, 2019; Osman et al., 2018). The 80 genes involved in the IPR pathway are controlled by *pals-22* and *pals-25* that are located next to each other on the genome. Together, *pals-22* and *pals-25* function as a molecular switch between growth and antiviral defense. The gene *pals-22* promotes development and lifespan, whereas *pals-25* stimulates pathogen resistance (Reddy et al., 2017, 2019). Of the 80 IPR genes that become differentially expressed upon infection, 25 genes belong to the *pals*-gene family. Although the function of most *pals*-proteins remains opaque, PALS-22 and PALS-25 physically interact together and are likely to interact with additional PALS-proteins (Panek et al., 2020). The total *pals*-gene family contains 39 members mostly found in five genetic clusters on chromosome I, III, and V (Chen et al., 2017; Leyva-Díaz et al., 2017). Both the antiviral IPR and the antiviral RNAi pathway require presence of *drh-1* that likely functions as a viral sensor (Ashe et al., 2013; Sowa et al., 2019).

At present, natural populations of *C. elegans* have been isolated worldwide and catalogued into 330 isotypes maintained by the *C. elegans* Natural Diversity Resource (CeNDR) (Cook et al., 2017). The collection contains *C. elegans* nematodes from every continent except Antarctica and each isotype in the CeNDR collection has been sequenced (Cook et al., 2017). Previous research found that genetic variations in the genes *drh-1* and *cul-6* change susceptibility to the OrV (Ashe et al., 2013; Sterken et al., 2021), yet most likely additional genetic variants can influence host-virus interactions in nature. The CeNDR database provides an ideal platform to investigate worldwide genetic variation and traces of evolutionary selection in antiviral genes in *C*. elegans.

Current studies investigating the IPR in *C. elegans* have focused on the reference genotype Bristol N2 (Reddy et al., 2017, 2019; Sowa et al., 2019; Panek et al., 2020). Here we set out to examine if the *pals*-genes experience selective pressure by analyzing the genetic variation in 330 wild strains from the CeNDR database. The *pals-*gene family is defined by the commonly shared ALS2CR12 domain and is expanded in *C. elegans* (humans and mice only contain a single *pals*-gene ortholog) (Leyva-Díaz et al., 2017). Expanded gene families often result from evolutionary selection (Thomas, 2006), suggesting that genetic variants in the *pals*-family could determine viral susceptibility. We found that only a few haplotypes occur worldwide for the *pals*-genes and that some are in regions of ancient genetic origin. This indicates that different pools of *pals-*genes have been maintained in *C. elegans* populations: a hallmark of balancing selection. Genetic variation in the *pals*-gene family, and specifically in the IPR-regulators, *pals-22* and *pals-25*, affects susceptibility to viral infection. This phenotype is further explored by infecting two well-studied strains, Bristol N2 and the Hawaiian strain CB4856 (Sterken et al., 2021), representing distinct *pals-22 pals-25* haplotypes. Our data illustrate that regulatory genetic variation can determine (basal) IPR gene expression, suggesting that natural genetic variation in IPR genes may influence host-pathogen interactions in wild *C. elegans* populations.

## Materials and methods

### Nematode strains and culturing

*C. elegans* strains N2 (Bristol),CB4856 (Hawaii), JU1580, WN2002, DL238, JU310, NIC2, ECA396, JU1400, QX1794, EG4725, MY2693, ERT54 and ERT71 were used in the experiments. The strains ERT54 (jyIs8[*pals-5p::*GFP, *myo-2p::*mCherry] X) and ERT71 (jyIs15[F26F2.1p::GFP; *myo-2*::mCherry]) were kind gifts from Emily Troemel (Bakowski et al., 2014; Reddy et al., 2017, 2019). The strains DL238, JU310, NIC2, ECA396, JU1400, QX1794, EG4725 and MY2693 were obtained from CeNDR (Cook et al., 2017). Strains were kept on 6-cm Nematode Growth Medium (NGM) dishes containing *Escherichia coli* strain OP50 as food source (Brenner, 1974). In maintenance culture the temperature was kept at 12°C and the standard growing temperature for experiments was 20°C. Fungal and bacterial infections were cleared by bleaching (Brenner, 1974). The strains were cleared of males prior to the experiments by selecting L2 larvae and placing them individually in a well in a 12-wells plate at 20°C. Thereafter, the populations were screened for male offspring after 3 days and only the 100% hermaphrodite populations were transferred to fresh 9-cm NGM dishes containing *E. coli* OP50 and grown until starved.

### Orsay virus infection assay in liquid

Orsay virus stocks were prepared according to the protocol described before (Félix et al., 2011). After bleaching, nematodes were infected using 20, 50, or 100 µL Orsay virus/500 µL infection solution as previously described (Sterken et al., 2014). Mock infections were performed by adding M9 buffer instead of Orsay virus stock (Brenner, 1974). For each strain the maximum viral load was determined (4 biological replicates). The maximum viral load is the highest viral load that can be obtained for a strain and is reached when increasing amounts of virus do not significantly affect the viral load anymore (t-test, p > 0.05). For N2 and JU1580 20 µL of OrV sufficed to reach the maximum viral load. For CB4856 at least 50 µL OrV was needed to maximize the viral load. Hence, using 50µL OrV/500 µL infection solution the maximum viral load was obtained for all three strains which was therefore used in subsequent experiments (Figure 1B). Two virus stocks were used for these experiments: one for the first four biological replicates and one for the remaining four replicates.

**Figure 1.**
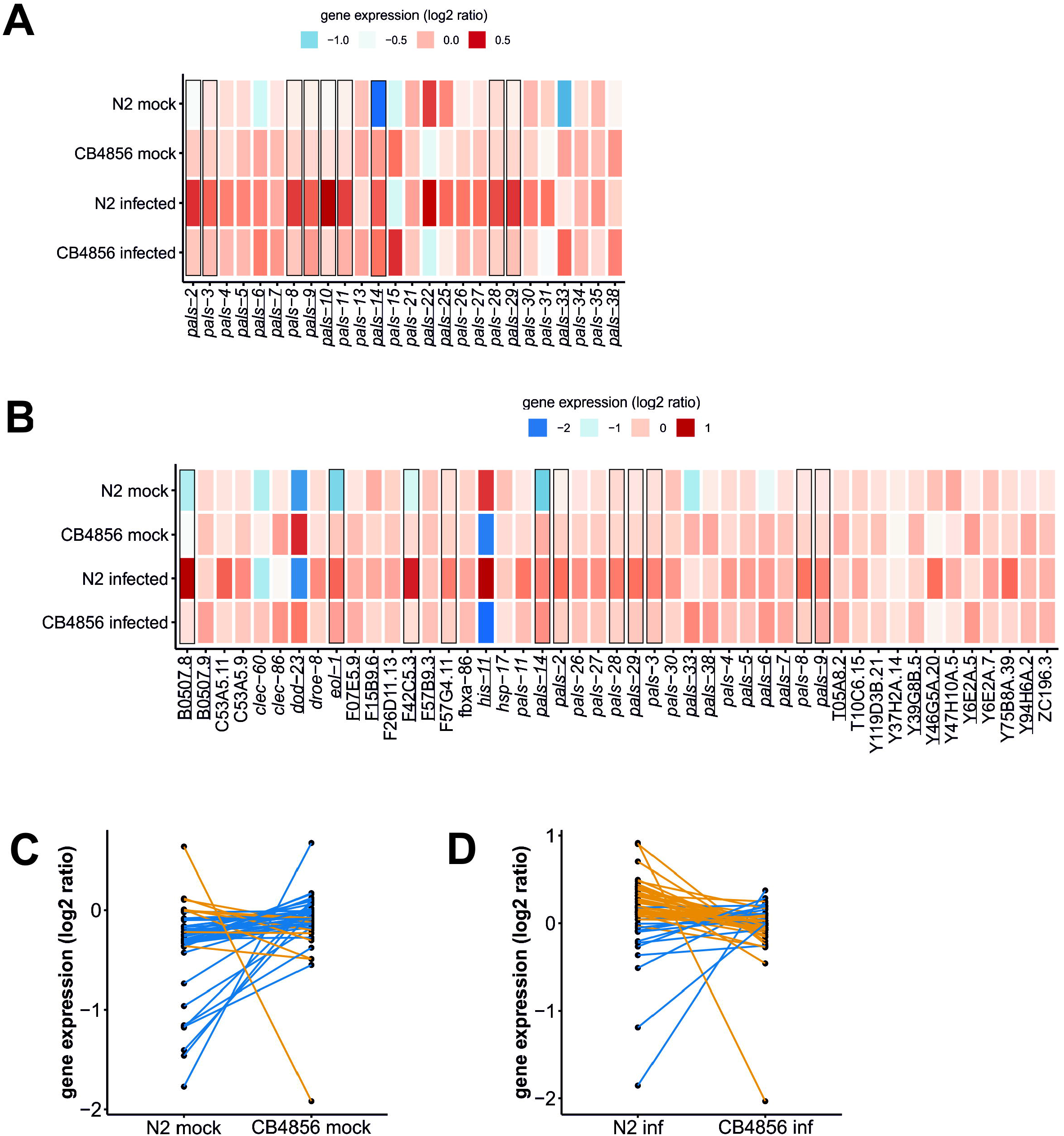
Natural variation in the C. elegans pals-gene family worldwide. A) The percentage of genetic variants (defined by SNPs) in the pals-gene family compared to the overall natural variation for each of the 330 wild isotypes (Cook et al., 2017). The number of SNPs is relative to the reference strain N2. Blue dots indicate the amount of variation in the pals-genes is different than expected from the overall natural variation (Chi-square test, FDR < 0.0001). B) Tajima’s D values per gene in the C. elegans genome calculated from sequence data of the 330 wild strains in the CeNDR database (Cook et al., 2017). Blue dots indicate Tajima’s D values for pals-genes.

The samples for the viral load and transcriptional analysis were infected in Eppendorf tubes with 50 µL Orsay virus/500µL infection solution 26 hours post bleaching (L2-stage) (8 biological replicates per treatment per genotype). The nematodes were collected 30 hours after infection. The samples for the transcriptional analysis of the time-series were infected with 50 µL Orsay virus/500 µL infection solution at 40 hours post bleaching (L3-stage). These strains were infected in the L3 stage to obtain high RNA concentrations for microarray analysis also in the early samples. The nematodes were collected at the following time points post-infection: 1.5, 2, 3, 8, 10, 12, 20.5, 22, 24, 28, 30.5, or 32 hours (1 biological replicate per treatment per genotype per time point). Viral loads of the samples were determined by RT-qPCR as described by (Sterken et al., 2014). A single Orsay virus stock was used for this experiment.

### Orsay virus infection assay on plate

The short-term plate exposure assay was used to infect 11 strains from 3 different *pals-22 pals-25* haplotypes. The protocol was adapted from previous experiments by (Chen et al., 2017; Reddy et al., 2017; Sowa et al., 2019) with OrV stock obtained as described by (Félix et al., 2011). Nematode populations were infected 22 hours post bleaching (L1 stage) and collected 24 hours after infection (L3 stage). Nematodes were infected with a mixture of 100µl OrV and 100 µl M9 solution that was spread equally over the plate. Before collecting the samples, the nematodes were washed three times with M9. Viral loads of the samples were determined by RT-qPCR as described previously (Sterken et al., 2014). RNA samples that had less than 25ng/µL RNA were excluded from the analysis. For this experiment a single OrV stock was used.

The long-term plate exposure assay was based on previous experiments by (Félix et al., 2011; Ashe et al., 2013) for which the Orsay virus stock was obtained as described previously (Félix et al., 2011). Three young adult N2 or CB4856 nematodes were placed on a plate with 20, 50, or 100 µL Orsay virus that was added to the plate shortly before transfer. M9 was added to mock-treated plates instead of Orsay virus stock. Two days after incubation part of the population was transferred to a fresh plate to prevent starvation. Four days (96 hours) after placing the first nematodes, populations were collected for RNA isolation. Viral loads of the samples were determined by RT-qPCR as described by (Sterken et al., 2014). A single Orsay virus stock was used for this experiment.

### Fluorescent in situ hybridization (FISH) of infected nematodes

Custom Stellaris FISH Probes were designed against OrV RNA1 by utilizing the Stellaris RNA FISH Probe Designer (Biosearch Technologies, Inc., Petaluma, CA) available online at www.biosearchtech.com/stellarisdesigner. The mock-treated or infected nematodes were hybridized with the Stellaris RNA FISH Probe set labeled with CAL Fluor® Red 590 Dye (Biosearch Technologies, Inc.), following the manufacturer’s instructions available online at www.biosearchtech.com/stellarisprotocols based on protocols by Raj *et al*. (Femino, 1998; Raj et al., 2008; Raj and Tyagi, 2010).

N2 and CB4856 populations were fixed 30h after infection (according to the short-term infection assay). Half of the nematodes were flash frozen to determine the viral load in the populations (Sterken et al., 2014) and the other half were used in the FISH procedure. Eight biological replicates were performed for this assay. The strains JU1580, ERT54, and ERT71 were mock-treated or OrV infected by chunking nematodes from a starved to a fresh plate and adding either 50 μL M9 or OrV. These nematodes were fixed for FISH 48 hours post mock-treatment or infection. For the reporter strains (ERT54 and ERT71) three biological replicates were performed and JU1580 nematodes were infected once. Nematodes were visualized using the Axio Observer Z1m inverted microscope (Zeiss).

### RNA isolation

The RNA of the samples in the transcriptional analysis (infected 26 hours post bleaching and collected 56 hours post bleaching) was isolated using Maxwell® 16 Tissue LEV Total RNA Purification Kit, Promega according to the manufacturer’s instructions including two modifications. First, 10 μL proteinase K was added to the samples (instead of 25 μL). Second, after the addition of proteinase K samples were incubated at 65°C for 10 minutes while shaking at 350 rpm. Quality and quantity of the RNA were measured using the NanoDrop-1000 spectrophotometer (Thermo Scientific, Wilmington DE, USA).

The RNA of the samples in the time series was isolated using the RNeasy Micro Kit from Qiagen (Hilden, Germany). The ‘Purification of Total RNA from Animal and Human Tissues’ protocol was followed, with a modified lysing procedure; frozen pellets were lysed in 150 µl RLT buffer, 295 µl RNAse-free water, 800 µg/ml proteinase K and 1% ß-mercaptoethanol. The suspension was incubated at 55°C at 1000 rpm in a Thermomixer (Eppendorf, Hamburg, Germany) for 30 minutes or until the sample was clear. After this step the manufacturer’s protocol was followed. Quality and quantity of the RNA were measured using the NanoDrop-1000 spectrophotometer (Thermo Scientific, Wilmington DE, USA) and RNA integrity was determined by agarose gel electrophoresis (3 μL of sample RNA on 1% agarose gel).

### cDNA synthesis, labelling and hybridization

The ‘Two-Color Microarray-Based Gene Expression Analysis; Low Input Quick Amp Labeling’ - protocol, version 6.0 from Agilent (Agilent Technologies, Santa Clara, CA, USA) was followed, starting from step five. The *C. elegans* (V2) Gene Expression Microarray 4×44K slides, manufactured by Agilent were used.

### Data extraction and normalization

The microarrays were scanned by an Agilent High Resolution C Scanner with the recommended settings. The data was extracted with Agilent Feature Extraction Software (version 10.7.1.1), following manufacturers’ guidelines. Normalization of the data was executed separately for the transcriptional response data (infected at 26 and collected at 56 hours post bleaching) and the transcriptional response of the time series. For normalization, R (version 4.0.2. x64) with the Limma package was used. The data was not background corrected before normalization (as recommended by (Zahurak et al., 2007)). Within-array normalization was done with the Loess method and between-array normalization was done with the Quantile method (Smyth and Speed, 2003). The obtained single channel normalized intensities were log_2_ transformed and the transcriptional response data (infected 26 hours post bleaching) was batch corrected for the two different virus stocks that were used for infection. The obtained (batch corrected) log_2_ intensities were used for further analysis using the package ‘tidyverse’ (1.2.1) in R (4.0.2, x64) (Wickham et al., 2019).

### Principal component analysis

A principal component analysis was conducted on the gene-expression data of the both the transcriptional response and the transcriptional response of the time series. For this purpose, the data was transformed to a log_2_ ratio with the mean, using

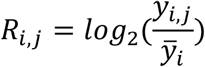

where R is the log_2_ relative expression of spot i (i = 1, 2, …, 45220) for sample j, and *y* is the intensity (not the log_2_-transformed intensity) of spot i for sample j. The principal component analyses were performed independently per experiment. The transformed data was used in a principal component analysis, where the first six axes that explain above 4.9% of the variance were further examined.

### Linear models

The log_2_ intensity data of the nematodes that were mock-treated 26 hours post bleaching and collected 56 hours post bleaching was analyzed using the linear model

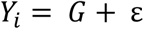

with Y being the log_2_ normalized intensity of spot i (1, 2, …, 45220). Y was explained over genotype (G; either N2 or CB4856) and the error term ε. The significance threshold was determined by the *p*.*adjust* function, using the Benjamini & Hochberg correction (FDR < 0.05) (Benjamini and Hochberg, 1995). The analyzed dataset is part of the dataset containing mock-treated and OrV infected samples.

The log_2_ intensity data of the nematodes that were either mock-treated or infected 26 hours post bleaching and collected 56 hours post bleaching was analyzed using the linear model

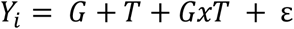

with Y being the log_2_ normalized intensity of spot i (1, 2, …, 45220). Y was explained over genotype (G; either N2 or CB4856), treatment (T, either infected or mock), the interaction between genotype and treatment and the error term ε. The significance threshold was determined by the *p*.*adjust* function, using the Benjamini & Hochberg correction (FDR < 0.1 for T and GxT, FDR < 0.05 for G) (Benjamini and Hochberg, 1995). Because of the minor effect of OrV infection on transcriptional activity, a relaxed false discovery rate (FDR < 0.1) was used to analyze the data. As all genes discovered using this threshold were IPR genes that are previously described by others, these were probably true positive hits (Sarkies et al., 2013; Chen et al., 2017)

The log_2_ intensity data for samples of the time series was analyzed using the linear model

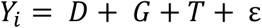

with Y being the log_2_ normalized intensity of spot i (1, 2, …, 45220). Y was explained over development (D, time of isolation: 1.5, 2, 3, 8, 10, 12, 20.5, 22, 24, 28, 30.5, or 32 hours post-infection), genotype (G; either N2 or CB4856), treatment (T; either infected or mock) and the error term ε. The significance threshold was determined by the *p*.*adjust* function, using the Benjamini & Hochberg correction (FDR < 0.05) (Benjamini and Hochberg, 1995). For the samples in the timeseries a correlation coefficient (r) was obtained by calculating the slope of gene expression over time.

The log_2_ intensity data of the nematodes that were exposed to heat shock (obtained from (Jovic et al., 2017)) was analyzed using the linear model

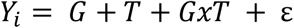

with Y being the log_2_ normalized intensity of spot i (1, 2, …, 45220). Y was explained over genotype (G; either N2 or CB4856), treatment (T, either control, heat shock or recovery), the interaction between genotype and treatment and the error term ε. The significance threshold was determined by the *p*.*adjust* function, using the Benjamini & Hochberg correction (FDR < 0.05) (Benjamini and Hochberg, 1995).

### Functional enrichment analysis

Gene group enrichment analyses were performed using a hypergeometric test and several databases with annotations. The databases used were: the WS258 gene class annotations, the WS258 GO-annotation, anatomy terms, phenotypes, RNAi phenotypes, developmental stage expression, and disease related genes (www.wormbase.org) (Stein et al., 2002; Lee et al., 2018); the MODENCODE release 32 transcription factor binding sites (www.modencode.org) (Gerstein et al., 2010), which were mapped to transcription start sites (as described by (Tepper et al., 2013)). Furthermore, a comparison with previously identified genes involved in OrV infection was made using a custom-made database (Supplementary Table S1).

Enrichments were selected based on the following criteria: size of the category n>3, size of the overlap n>2. The overlap was tested using a hypergeometric test, of which the p-values were corrected for multiple testing using Bonferroni correction (as provided by p.adjust in R, 4.0.2, x64). Enrichments were calculated based on unique gene names, not on spots.

### Probe alignment

Probe sequences of the *pals*-genes and IPR-genes (*C. elegans* (V2) Gene Expression Microarray 4×44K slides, Agilent) were aligned to the genome sequence of CB4856 (PRJNA275000) using command-line BLAST, using blastn with standard settings (Blast command line application; v2.2.28) (Altschup et al., 1990; Thompson et al., 2015; Cook et al., 2017). We also compared the *pals*-genes probes to a differential hybridization experiment, to see if differences in DNA sequence explain the mRNA-based hybridization differences. Therefore, we obtained data from Volkers *et al*., 2013 (normalized data, E-MTAB-8126) and corelated the gene-expression differences with the hybridization differences (Volkers et al., 2013).

### Gene expression measurements by RT-qPCR

Gene expression measurements were performed on the cDNA of each of the 32 samples used in the N2 and CB4856 gene expression analysis of 30 hours of mock-treated or infection (8 biological replicates) and on the cDNA of N2 and CB4856 mock-treated or infected samples exposed to 50µL on plate (4 biological replicates). Gene expression was quantified by RT-qPCR using custom designed primers (*pals-6* forward 5’-TGGGTTCTGGATCAAGCAAAT-3’, *pals-6* reverse 5’-TGTTCTAGAGCTGCCTGTCTCTG-3’, *pals-14* forward 5’-TCGGGAAAGCATCAATGAACTGC-3’, *pals-14* reverse 5’-TGTTGTGCCTCTCCTCTGCC-3’, *pals-22* forward 5’-TTTTAATCTTGAAAGTGACCGCTGGG-3’, *pals-22* reverse 5’-ACTCTCTGTTGTCGTCTTGCAAAATT-3’, *pals-25* forward 5’-TGCAATCCGAAGATTGGTGA-3’, *pals-25* reverse 5’-AAATTCTAACTTGCTCAGCATGGA-3’) that overlap at least one exon-exon border to prevent unintended amplification of any remaining DNA. RT-qPCR was performed on the MyIQ using iQ SYBR Green Supermix (Biorad) and the recommended protocol. Gene expression in each sample was quantified for the gene of interest and two reference genes (Y37E3.8 and *rpl-6*) (Sterken et al., 2014) in duplo.

To determine the relative gene expression, we normalized the data as in (Sterken et al., 2014). In short, we normalized the *pals*-gene expression based on the two reference genes using

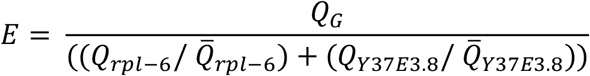

where E is the normalized gene expression, Q_G_ is the expression of the gene of interest, Q_rpl-6_ is the expression of the reference gene *rpl-6* and Q_Y37E3.8_ is the expression of the reference gene Y37E3.8.

### Genetic variation analysis

Genetic data on *C. elegans* wild strains were obtained from the CeDNR website (release 20180527) (Cook et al., 2017). The data was further processed using custom made scripts (https://git.wur.nl/mark_sterken/Orsay_transcriptomics). In short, the number of polymorphisms in the *pals-*family within a strain was compared to the total number of natural polymorphisms found in that that strain. The N2 strain was used as the reference strain. A chi-square test (FDR < 0.0001) was used to determine whether strains showed less or more variation than expected within the *pals*-gene family compared the total natural variation observed. Next, we also manually inspected the *pals-22 pals-25* locus of each of the 330 isolates via the Variant Browser tool on the CeNDR website (www.elegansvariation.org) (Cook et al., 2017). The *pals-22 pals-25* locus could be classified in three major groups based on structural variation observed in the bam-files.

The number of polymorphisms within the *pals*-gene family was further specified per gene. Tajima’s D values were calculated per gene within the *C. elegans* genome using the PoPGenome package (Pfeifer et al., 2014). The number of polymorphisms within the *pals*-gene family were compared to the geographical origin of the strain obtained from the CeDNR database (Cook et al., 2017). The data were visualized using the packages ‘maps’ (3.3.0) and ‘rworldmap’ (1.3-6) (Becker and Wilks, 1993, 1995; South, 2011; Brownrigg et al., 2018).

### eQTL data analysis

The eQTL data was mined from https://bioinformatics.nl/EleQTL (Snoek et al., 2020).

### Data analysis and availability

All custom written scripts were made in R (4.0.2, x64) and the script and underlying data are available via https://git.wageningenur.nl/published_papers/sluijs_etal_2021_orv_transcriptomics. The transcriptome datasets generated are deposited at ArrayExpress (E-MTAB-7573 and E-MTAB-7574). The data of the 12 N2 mock samples of the time series has previously been described (Snoek et al., 2015).

## Results

### Global genetic variation in the pals-family is shaped by balancing selection

The Intracellular Pathogen Response (IPR) counteracts viral infection in *Caenorhabditis elegans* and involves activity of 25 *pals*-genes (Reddy et al., 2017, 2019). To examine genetic variation in the *pals-* family genes, we investigated sequence information from the 330 wild strains from the CeNDR database (Cook et al., 2017). For each wild strain the genetic variation (compared to the reference strain N2) was summarized for genes in the *pals*-family and for all genes. For 48 wild strains genetic variation (defined by SNPs) in the *pals*-family was higher than expected compared to the overall genetic variation (chi-square test, FDR < 0.0001), but for 204 strains of the 330 analyzed strains the *pals-*gene family contained less variation than the overall genetic variation (chi-square test, FDR < 0.0001) (Figure 1A) (see Material and Methods for details). This indicated that while the *pals-*genes belong to an expanded gene family, most wild strains contain relatively little genetic variation in the *pals-*genes compared to N2.

Populations from distinct geographical locations may encounter different selective pressures (Sivasundar and Hey, 2005; Volkers et al., 2013). However, after mapping the amount of natural variation to the geographical location, no clear geographical pattern could be found (Supplementary Figure S1). Interestingly, some local strains show highly diverging levels of genetic diversity within the *pals*-family. For example, strain WN2002 was isolated in Wageningen (the Netherlands) and contains three times more genetic variation in the *pals*-family than the average of other genes. Strain WN2066 was isolated from the same compost heap as WN2002. Yet, compared to N2, WN2066 has high conservation of the *pals*-genes (0.27% SNPs), despite higher overall genetic variation (2.67% SNPs). This shows that at the same geographic location, genetic diversity in the locus can be retained in the population, possibly due to differential microenvironmental pressures.

Next, we tested whether DNA sequence divergence was subjected to genetic drift, or that selective forces were acting on the *pals*-family. Overall Tajima’s *D* (TD) values in *C. elegans* populations are low as a result of overall low genetic diversity (TD_mean_ = −1.08, TD_median_ = −1.12) (Andersen et al., 2012), but four *pals*-genes (*pals-17, pals-18, pals-19*, and *pals-30*) showed positive TD values suggesting either balancing selection or a low frequency of rare alleles (Figure 1B). The most clear example was *pals-30* that had a TD value of 4.8: the highest value of all tested *C. elegans* genes. In total, 11 out of 39 *pals-*genes had values that fall within the 10% highest TD values for *C. elegans* (TD > −0.42) and these top 10% genes included IPR regulators *pals-22* and *pals-25* (Supplementary Table S2).

Subsequently, we delved into the genetic diversity for each *pals*-gene by investigating the number of variants per *pals*-gene (Supplementary Figure S2, Supplementary Table S2). Several *pals*-genes contained hardly any genetic variation and were therefore conserved on a worldwide scale. This conserved group contains the gene *pals-5* which acts downstream in the IPR (Reddy et al., 2017). Other *pals*-genes were highly variable, sometimes containing hundreds of polymorphisms (SNPs) in a singlegene. Interestingly, for most genes in the diverse group, few alleles exist worldwide. For example, three alleles were found for the gene *pals-25*: strains that harbor an N2-like allele, an allele containing ∼30 polymorphisms (the well-studied Hawaiian strain CB4856 belongs to this group) or an allele containing ∼95 polymorphisms (illustrated by the strain WN2002 from the Netherlands). In total, 19 out of 24 highly variable *pals*-genes show a clear grouping within two or three haplotypes suggesting that these haplotypes are actively maintained in the populations which supports that balancing selection could be acting on these genes (Supplementary Figure S2, Supplementary Table S2).

Manual inspection of the mapped reads in the 330 CeNDR strains showed evidence for extensive polymorphisms in the *pals-22 pals-25* locus that regulate the IPR transcriptional response (Reddy et al., 2017, 2019). In total, we found three major *pals-22 pals-25* haplotypes (N2-like, CB4856-like, and WN2002-like) that occur globally (Figure 2A-D, Supplementary Table S3) with the highest local genetic diversity found on the Hawaiian islands and Pacific region (Cook et al., 2017; Crombie et al., 2019; Lee et al., 2021). The genetic variation in the region where *pals-22* and *pals-25* are located is estimated to have diverged 10^6^ generations ago (Thompson et al., 2015; Lee et al., 2021). Notably, *pals-22* and/or *pals-25* are both thought to have early stop codons in CB4856 and WN2002 that could change or disrupt their functioning, in particular in *pals-25*, where the stop codon is located before the ALS2CR12 domain (Cook et al., 2017; Leyva-Díaz et al., 2017). Yet, poor mapping to the reference genome in these highly diverse regions hampers the reliability of exact variant calling; in 24 wild strains most of the intron sequence was not covered by reads at all. The latter suggests that the genetic sequence in those strains is highly polymorphic and additional in-depth sequencing of the strains would be necessary to fully uncover the genomic sequence at these locations. In conclusion, within the *pals-*gene family we found genes with either globally conserved or a few genetically distinct alleles. In particular, the *pals*-genes with a division into a few haplotypes show atypically high Tajima’s D values when compared to other *C. elegans* genes. Together, our findings indicate that the *pals*-genes have been experiencing evolutionary pressure that resulted in long-term balancing selection of these genes.

**Figure 2.**
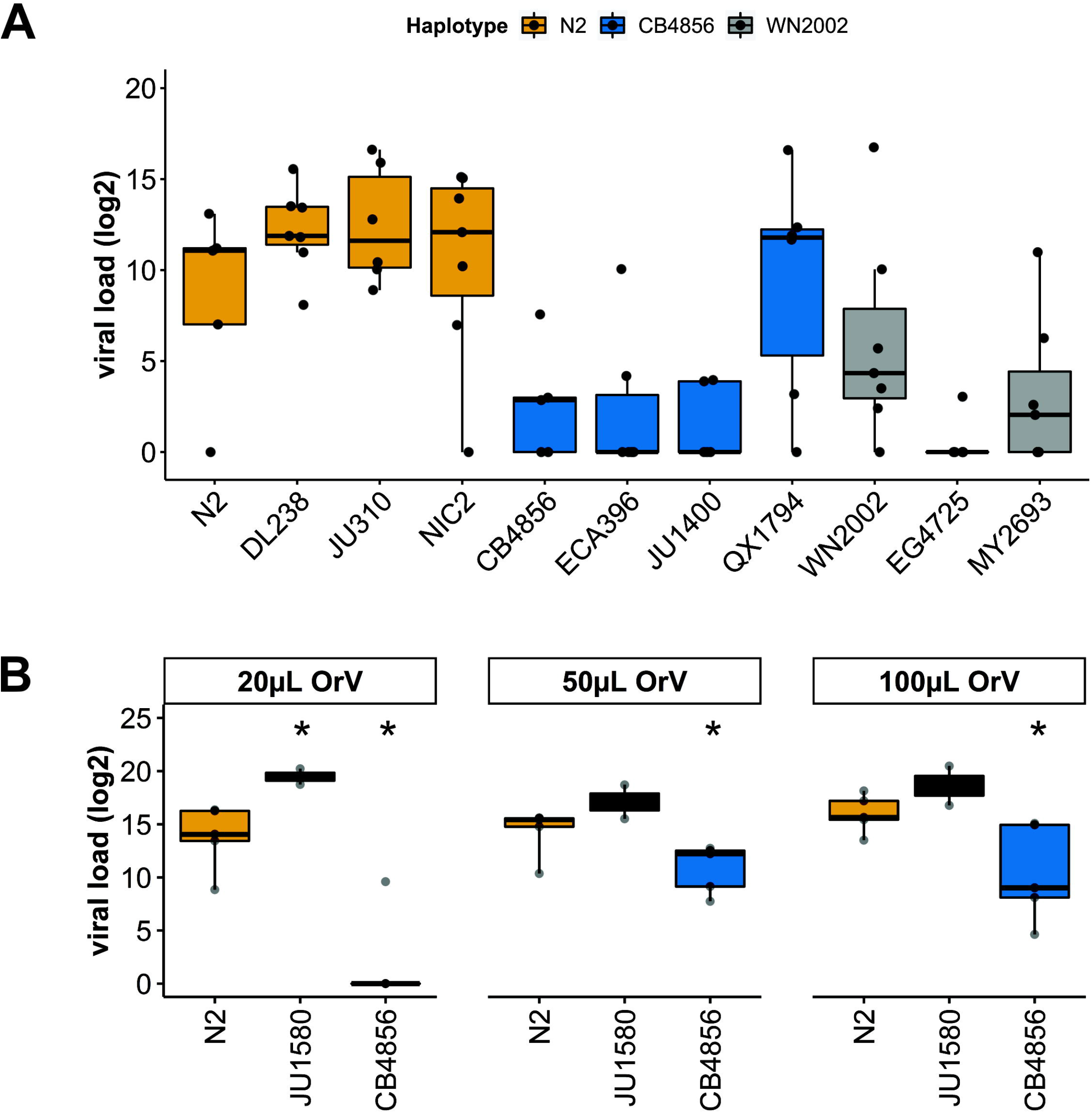
Worldwide haplotype diversity found for pals-22 and pals-25. A) Three distinct haplotypes were found at the pals-22 pals-25 locus, here represented by an illustration of the read coverage at the locus. The most common haplotype (N2-like) is found in 269 stains and shows low coverage of the second intron of pals-22. The second common haplotype is CB4856-like and was found in 31 strains. For these strains coverage indicates strong structural variation in the introns of both genes, as well as larger insertions/deletions. Then, the WN2002-like haplotype was found in 28 strains and consists of very extensive structural variation at the locus, where almost the entire intron structure is not covered by reads. B) A geographical representation of the pals-22 pals-25 haplotypes found worldwide. C) Zoomed in representation of Figure S2B of the strains collected in Europe. D) Zoomed in representation of Figure S2B of the strains collected on Hawaii.

### The antiviral response in strains with distinct pals-22 pals-25 haplotypes

To investigate if the *pals-22 pals-25* haplotype determined viral susceptibility of strains, eleven genetically divergent *C. elegans* strains from the CeNDR collection were infected with OrV. These eleven strains (N2, DL238, JU310, NIC2, CB4856, ECA396, JU1400, QX1794, WN2002, EG4725 and MY2693) all contain a *drh-1* allele without deletions (Cook et al., 2017), because an intact *drh-1* allele is essential for antiviral IPR activation (Sowa et al., 2019). Nematodes were infected in the L1-stage and exposed to viral infection for 24-hours after exposure. The *pals-22 pals-25* haplotype explained viral susceptibility in the dataset (Figure 3A) (37% variance, p = 5·10^−6^), besides the individual genotype of the strain (Figure 3A) (17% variance, p = 0.02). In general, strains with a N2 *pals-22 pals-25* haplotype were more susceptible to OrV than strains with a CB4856 or WN2002 *pals-22 pals-25* haplotype. Furthermore, OrV was only able to reproduce in 55% of infected samples with a CB4856 or WN2002 haplotype, compared to 92% of the samples with an N2 haplotype. Therefore, extensive genetic variation within the *pals-22 pals-25* locus (compared to the N2 reference) enhanced resistance to viral infection.

**Figure 3.**
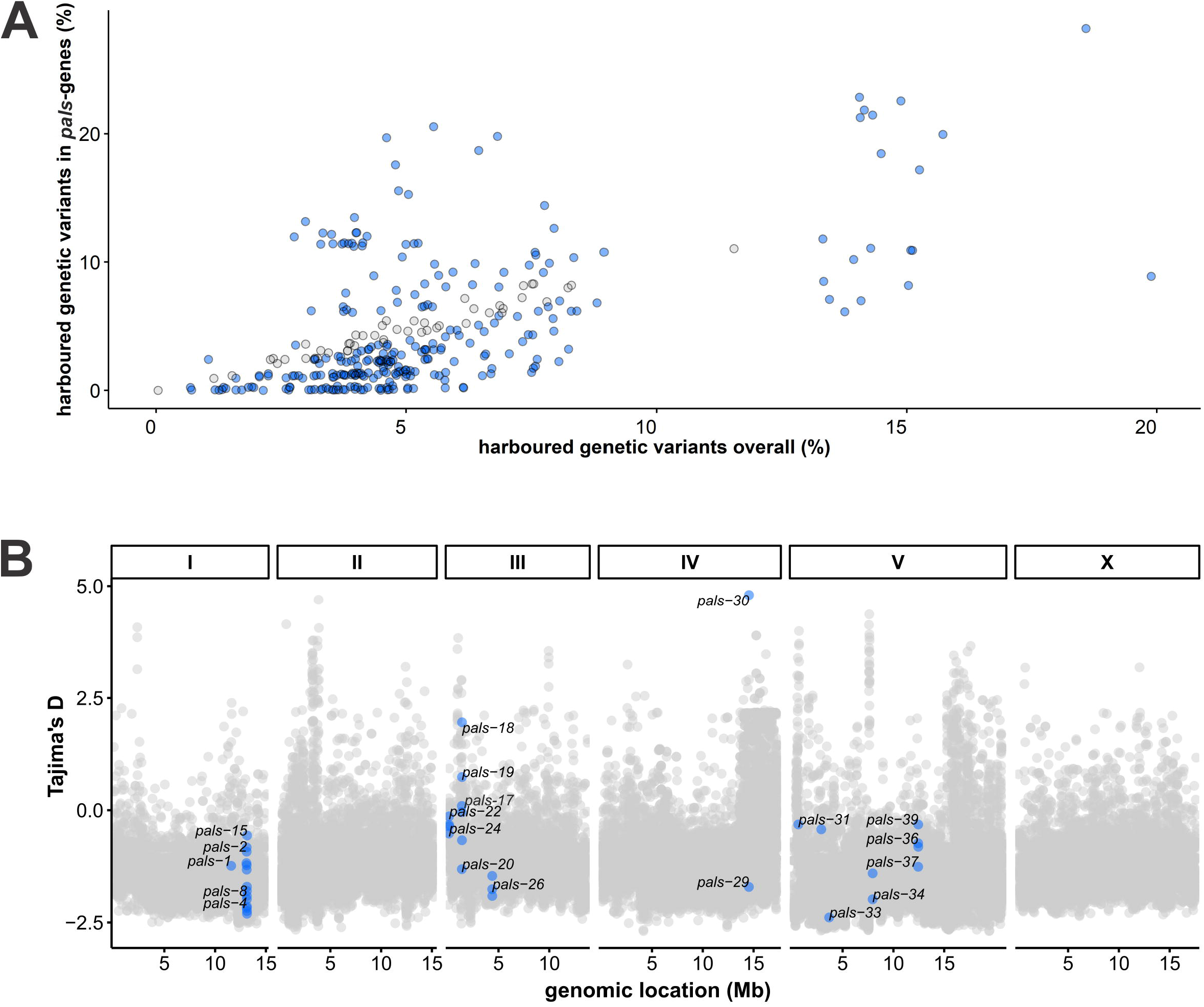
Viral susceptibility of C. elegans N2, CB4856 and JU1580 upon initial infection. A) Viral loads (log_2_) as determined by RT-qPCR for three different haplotypes after exposure to 100µL OrV on the plate. The N2 haplotype (N2, DL238, JU310, NIC2 strains) is shown in orange, the CB4856 haplotype (CB4856, ECA396, JU1400, QX1794 strains) is shown in blue and the WN2002 haplotype (WN2002, EG4725 and MY2693) is shown in grey. B) Viral loads (log_2_) as determined by RT-qPCR for the strains N2, CB4856 and JU1580 after exposure to 20, 50 or 100µL OrV/500µL infection solution (student t-test; * p < 0.05).

To further study the antiviral response for distinct *pals-22 pals-25* haplotypes, we compared the strains N2 and CB4856 in more detail. N2 and CB4856 are well-characterized genotypes that differ in viral susceptibility to the Orsay virus (Thompson et al., 2015; Sterken et al., 2021). This difference in viral susceptibility could be partially explained by polymorphisms in the antiviral gene *cul-6* as revealed by a linkage mapping but the majority of the variation in the phenotype remained unexplained (Sterken et al., 2021). Here, N2 and CB4856 nematodes were exposed to varying concentrations of the OrV (Sterken et al., 2014, 2021). Additionally, JU1580 nematodes were taken along as a highly susceptible control (Félix et al., 2011; Sterken et al., 2014). We confirmed that CB4856 was less susceptible than N2 after exposure to different concentrations or OrV and that JU1580 was more susceptible than N2 (Figure 3B) (Félix et al., 2011; Sterken et al., 2014, 2021). Moreover, we explored the difference in the N2 and CB4856 phenotype further by staining infected nematodes using Fluorescent *in situ* Hybridization (FISH). We found that the OrV could only be detected in a minor faction of nematodes (<1%) precluding a direct quantitative comparison between these two strains. Therefore, the infection does neither reach high levels of infection in N2 nor CB4856 30h post infection (Supplementary Table S4). Still, FISH staining of IPR gene reporter strains ERT54 and ERT71 showed that low levels of OrV can already activate the IPR (Supplementary text S1, Figure S3 and Supplementary Table S3), indicating that use of FISH could underestimate the number of animals responding to infection. Therefore, we continued with measuring IPR expression in infected N2 and CB4856 nematodes.

### Basal IPR expression differs between N2 and CB4856

Distinct *pals-22 pals-25* haplotypes may result in distinct IPR activity between wild strains. To study whether the N2 and CB4856 *pals-22 pals-25* haplotypes underlie differential IPR gene expression, we measured their transcriptomes using microarrays under standard conditions and after exposure to the OrV (Figure 3B). Although microarrays were originally designed for the N2 strain, they have been used and tested repeatedly for the CB4856 strain (Capra et al., 2008; Rockman et al., 2010; Viñuela et al., 2012; Volkers et al., 2013; Thompson et al., 2015; Snoek et al., 2017). Based on these studies, we have identified 17 IPR- and 11 *pals-*gene probes with incorrect alignment to the CB4856 genome and excluded these from any further analyses (Supplementary Table S5A-E).

First, we focused on transcriptional differences between the N2 and CB4856 strain in the mock-experiment. Expression patterns of the full dataset were analyzed by means of a principal component analysis (PCA). Genotype explained the main difference in gene expression patterns (36.1%), which is in line with previous results, see for example (Li et al., 2006; Capra et al., 2008; Volkers et al., 2013; Snoek et al., 2017) (Supplementary Figure S4). Among the 6383 genes (represented by 9379 spots) that were differentially expressed between N2 and CB4856 (under mock conditions) were 131 genes known to be involved in OrV infection (Supplementary Table S1, S6A). These include twenty-three IPR genes – including ten *pals*-genes – showing significantly higher expression in CB4856 compared to N2 (Figure 4A, B) (FDR < 0.05). In general, most IPR- and *pals*-genes appeared more active in CB4856 (Figure 3C). Expression levels measured by RT-qPCR confirmed the microarray data for *pals-6, pals-14*, and *pals-22*, although the slight difference in *pals-25* expression found on the microarrays was not replicated (Supplementary Figure S5). Contrary to most other *pals-*genes, *pals-22* expression is higher in N2 than in CB4856 (Figure 4A) (FDR < 0.05, Supplementary Table S6A) (Vu et al., 2015), which may determine gene expression of other IPR members (Figure 4B) (Reddy et al., 2019). Concluding, the IPR was overall more active in CB4856 than in N2 under standard conditions.

**Figure 4.**
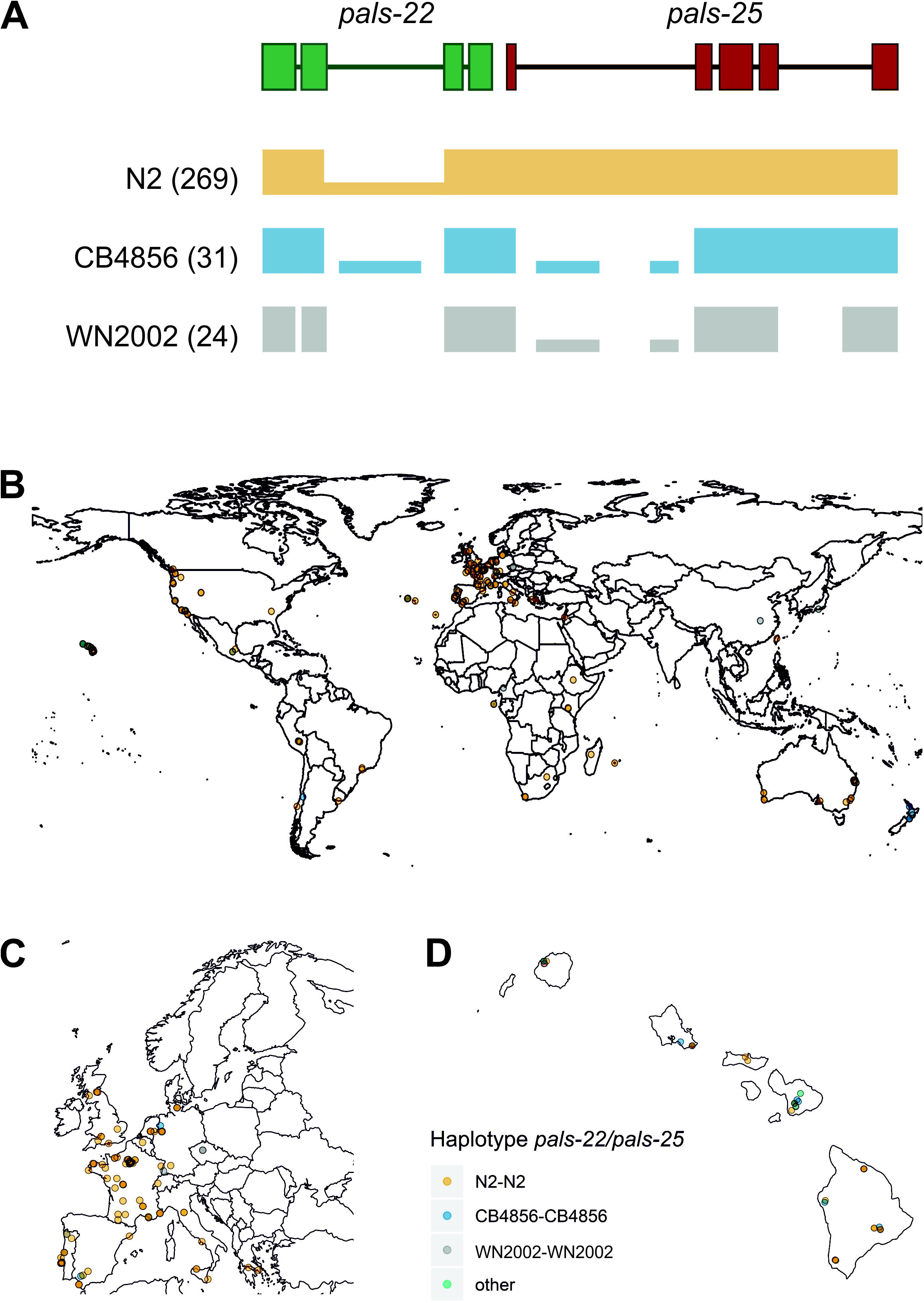
Gene expression of IPR and pals-genes in C. elegans N2 and CB4856 under control and OrV-infected conditions. A) Heat-map showing the log_2_ intensities of pals-genes in N2 mock, N2 infected, CB4856 mock and CB4856 infected conditions B) Heatmap showing the expression of IPR genes in log_2_ intensities in N2 mock, N2 infected, CB4856 mock and CB4856 infected conditions. Underlined genes showed significant (basal) expression differences based on genotype (FDR < 0.05), whereas squares indicated the genes where treatment- or the combination of treatment and genotype had a significant effect (FDR < 0.1) (Supplementary Table S6). Log_2_ ratios are based on the average expression of the gene of interest in the overall dataset. Therefore, the log_2_ ratios per experimental group indicate the deviation from the average value. Please note, a subset of the pals-genes, namely the pals-genes that are also IPR genes (defined in (Reddy et al., 2019)) are depicted twice. This allows for direct comparison to other pals-genes that do not become differentially expressed upon infection (like pals-22 and pals-25) and to non-pals IPR genes. C) Per IPR gene comparison of the log_2_ ratios in N2 and CB4856 samples under mock conditions. Blue lines show genes that on average showed higher expression in CB4856 (38 genes), orange lines connect genes that on average showed higher expression in N2 (9 genes). D) Per IPR gene comparison of the log_2_ ratios in N2 and CB4856 samples under infected conditions. Blue lines show genes that on average showed higher expression in CB4856 (14 genes), orange lines connect genes that on average showed higher expression in N2 (33 genes).

Next, we analyzed the transcriptomes of nematodes collected 30 hours post infection (Figure 4). As expected, based on a PCA where samples did not separate based on infection status, relatively few genes responded to the OrV in our experiment (Supplementary Figure S4, Supplementary Table S6B) (Sarkies et al., 2013; Chen et al., 2017). Gene expression analysis by a linear model showed that 27 genes (represented by 57 spots) were differentially expressed upon infection by OrV (FDR < 0.1) (Supplementary Figure S6A) and 18 genes (represented by 44 spots) were differentially expressed by a combination of both treatment and genotype (FDR < 0.1) (Supplementary Figure S6B). These two groups of genes were largely overlapping (Supplementary Figure S6C) and most of these genes only respond to infection in the genotype N2 (Figure 4B). Many of the *pals*-gene family members became higher expressed after infection in N2, but not in the strain CB4856 (Figure 4A). This led to a slightly more active IPR in N2 under infected conditions than in CB4856 (Figure 4D).

Thus, N2 showed a IPR to OrV infection, however we did not detect increased expression of IPR genes in CB4856 nematodes. Yet, the IPR can also be activated by heat stress and by re-analyzing a previous dataset we observed that *pals-*genes were activated after heat shock in CB4856 (Supplemental Text S2, Supplemental Figure S7) (Jovic et al., 2017, 2019). Therefore, the lack of a transcriptional response 30h post OrV infection could result from early activation of IPR genes or the infection stress being too mild stress to trigger the IPR. We tested the first hypothesis by measuring IPR activity over a 30-hour time-course. This did not show evidence for an earlier IPR in CB4856 than N2 although we noticed that OrV responsive genes in CB4856 were more dynamic (they showed more fluctuation in expression) under both standard and infected conditions than in N2 (Figure S8A, Supplemental Table S6C). The second hypothesis was tested by exposing N2 and CB4856 continuously to OrV for four days. A previous study indicated that viral loads in these mixed-staged populations were comparable between N2 and CB4856 and we hypothesized long-term exposure would therefore lead to higher viral pressure (Ashe et al., 2013). We first confirmed the previously found similarity between viral load in N2 and CB4856 four days after exposure after which both strains had higher viral loads than than after 30h of exposure (Figure S9). Subsequently, gene expression of *pals-6, pals-14, pals-22*, and *pals-25*, was measured for long-term infected N2 and CB4856 populations. We found that *pals-6* and *pals-14* were upregulated in CB4856 (Figure S8B). Together, these experiments indicate that sufficiently high stress can raise IPR expression levels in CB4856.

## Discussion

Viral susceptibility can be determined by host genetic variation. Here, we have studied the effect of host genetic diversity on natural viral infection in the nematode *C. elegans*. Our findings show that genetic variation in *C. elegans* affects the Intracellular Pathogen Response (IPR): a transcriptional response that counteracts pathogens by increased proteostasis and in which at least 27 *pals-*genes are involved (Reddy et al., 2017, 2019). The 39 members of the expanded *pals-*gene family are mostly conserved within the *C. elegans* species and the *pals-*genes for which divergent alleles do occur can be clustered into a few different haplotypes. We found that *pals-22 pals-25* haplotype determines the viral susceptibility of genetically highly distinct strains. Furthermore, studying the transcriptome of two strains with different *pals-22 pals-25* haplotypes showed that genetic variation directs basal IPR activity. Therefore, this study reveals natural variation in the IPR that protects wild *C. elegans* from viral, oomycete and microsporidian infection.

### IPR genes of the pals-family are under balancing selection

Population genetic analyses showed that IPR genes are experiencing selective pressure which could be a result of balancing selection, population bottlenecks or presence of rare genetic variants. We argue that balancing selection is the most likely cause for three reasons. First, *pals-22* and *pals-25* were experimentally validated to regulate the IPR and to balance growth and immunity (Reddy et al., 2019). Second, we observed that few major haplotypes occur for this gene-pair and most other *pals-*genes. Manual inspection of the *pals-22 pals-25* locus did not suggest presence of rare variants, rather the presence of a highly divergent region of ancient origin (Thompson et al., 2015). Third, presence of the *pals-22 pals-25* divergent region did not correlate with overall genetic variation, hence is unlikely to be the result of a bottleneck.

Besides *pals-22* and *pals-25*, multiple other *pals*-genes studied here show signs of balancing selection (high Tajima’s D values) (Tajima, 1989), in particular the genes on the first and second cluster on chromosome III (0.1 and 1.4Mb). In contrast, most of the genes in *C. elegans* show negative Tajima’s D values due to a recent selective sweep affecting chromosome I, IV, V, and X. This selective sweep greatly reduced the genetic variation within the species (Andersen et al., 2012). The *pals*-genes with relatively high Tajima’s D values on chromosome III are located in a region that has diverged early in the natural history of *C. elegans* (Thompson et al., 2015; Lee et al., 2021). Despite this ancient divergence, few haplotypes occur for this region and only a minority of strains, including CB4856, carry genetic variants distinct from N2.

Genetic variation in the *pals*-gene family regulates an evolutionary important transcriptional response to environmental stress, which includes pathogens. Given the minor effect of OrV on fecundity (Félix et al., 2011; Ashe et al., 2013), it seems unlikely that the OrV is one of the pathogens that exert selection pressure underlying the balancing selection. However, immunity responses upon microsporidia and oomycete infection are also mediated by the IPR (Bakowski et al., 2014; Reddy et al., 2017, 2019; Osman et al., 2018). As these pathogens are lethal (Zhang et al., 2016), we think that it is possible that these classes of pathogens underlie maintenance of different IPR haplotypes in natural populations. A recent example shows balancing selection in the plant genus *Capsella* also results in maintenance of ancestral genetic variation in immunity genes. The two *Capsella* species studied retained genetic variably at immunity loci, despite a recent population bottleneck and reproduction by selfing that together reduced overall genetic variation. Here, parallels can be drawn to *C. elegans*, a species that also mainly reproduces by selfing and has experienced loss of global genetic diversity (Andersen et al., 2012). Together, these studies show that within natural populations immunity-related genetic variation can be retained by balancing selection.

### Transcriptional activation of the IPR in genetically diverse strains

CB4856 shows a high basal expression of multiple IPR genes which may be possible due to regulatory genetic variation in the *pals*-genes. Most of the genetically diverse *pals*-genes on chromosome III and V have previously been shown to display local regulation of gene expression in N2xCB4856 recombinant inbred lines (*cis*-quantitative trait locus; *cis-*eQTL) (Snoek et al., 2020). Moreover, at least 10 genes across different *pals*-clusters were regulated by genes elsewhere in the genome (*trans*-eQTL) (Snoek et al., 2020). Most of these expression QTL were consistently found across multiple studies, environmental conditions and labs (Li et al., 2006, 2010; Rockman et al., 2010; Viñuela et al., 2010, 2012; Sterken et al., 2014; Snoek et al., 2017). The established IPR regulators *pals-22* and *pals-25* could be likely candidates for this regulatory role.

Although our data demonstrates that the strain CB4856 has multiple IPR genes with higher basal expression than in the strain N2 it remains unclear whether this leads to lower susceptibility to OrV infection. We observe lower viral RNA accumulation in CB4856 compared to N2 during the first 30 hours of infection, therefore high basal IPR expression may slow the infection. During this initial period of viral infection, we did not detect upregulation of IPR genes in CB4856 compared to its basal expression. Yet, after a longer period of viral exposure on the plate, CB4856 accumulates as much virus in the population as N2 and subsequently some IPR genes were also upregulated in CB4856. Therefore, plate infection assays may evoke stronger transcriptional responses which could explain why previous studies found more differentially expressed genes than this study (Sarkies et al., 2013; Chen et al., 2017). Furthermore, transcriptional techniques, such as single-cell RNA-seq or TOMO-seq (Trapnell et al., 2017; Ebbing et al., 2018), may provide more details about the local transcriptional response within infected cells and studying gene expression in additional strains could demonstrate the generality of basal IPR expression in relation to the *pals-22 pals-25* IPR haplotype. and

### Are there alternative IPR strategies?

Strains potentially harbor regulatory genetic variation tailored to specific environments. In a harsh environment constant activity of the IPR may be preferred over low expression. Finding out which environmental factor could explain the population genetic patterns within the *pals*-genes of the IPR will be challenging. The IPR pathway has been shown to respond to multiple environmental stressors including intestinal and epidermal pathogens, but also heat stress (Reddy et al., 2017, 2019). Despite the increasing amount of ecological data for both *C. elegans* (Cook et al., 2017) and its pathogens (Zhang et al., 2016; Richaud et al., 2018; Frézal et al., 2019), it is not yet sufficient to draw any firm conclusions whether co-occurrence of host and pathogen drives evolution within the *pals*-family. However, some evidence exists that host-pathogen interactions can affect the genotypic diversity at a population level. In Orsay (France), the location where OrV is found, diversity in pathogen susceptibility potentially explains the maintenance of several minority genotypes. These minority genotypes are outcompeted in the absence of the intracellular pathogen *Nematocida parisii*, but perform better in the presence of the pathogen (Richaud et al., 2018). Perhaps this can also help explain our observations of divergent *pals-22 pals-25* haplotypes in strains found at the same site. Experimental evolution experiments hold the potential to bridge this gap between the lab and the field by investigating if the presence of intracellular pathogens invokes any genetic and transcriptional changes within the *pals*-family (Gray and Cutter, 2014; Teotonio et al., 2017).

Taken together, this study provides insights into the natural context of the evolutionary conserved genetic and the plastic, transcriptional response after infection. We show that relatively little genetic diversity is found worldwide within clusters of *pals-*genes that regulate the IPR transcriptional response. In addition, the genetic diversity that exists is captured by only a few highly divergent haplotypes occurring worldwide. Therefore, we suggest that genes that function in the IPR transcriptional response could be under balancing selection, possibly from intracellular pathogens. Our results show the haplotype of the IPR regulators *pals-22* and *pals-25* determined the viral susceptibility of genetically distinct strains and that genetic variation within wild *C. elegans* can shape the basal expression of IPR genes. Thereby, this study provides new insights into the diversity of ways that hosts can develop both genetic and transcriptional responses to protect themselves from harmful infections.

## Author contributions

BLS, GPP, JEK and MGS conceived and designed the experiments. LvS, KB, FP, TB, JIHAW, JAGR, and MGS conducted the experiments. LvS and MGS conducted transcriptome and main analyses. LvS, GPP, JEK, and MGS wrote the manuscript. All authors read and provided comments on the manuscript.

## Acknowledgements

The authors want to thank Erik Andersen for hosting and sharing natural variation data on CeNDR and his advice on population genetic analyses. Marie-Anne Félix is kindly thanked for sharing the Orsay virus and Emily Troemel for sharing IPR reporter strains.

## Data availability

Strains used can be requested from the authors. The transcriptome datasets generated are deposited at ArrayExpress (E-MTAB-7573 and E-MTAB-7574). All custom written scripts and data for the analyses in R can be found at https://git.wageningenur.nl/published_papers/sluijs_etal_2021_orv_transcriptomics.

