## Supplementary for "Balancing selection of the Intracellular Pathogen Response in natural *Caenorhabditis elegans* populations"

**Supplementary Text S1.** ***Fluorescent in situ Hybridization of OrV reporter strain nematodes***

Using Fluorescent *in situ* Hybridization (FISH) staining of OrV we could only visualize a few infected cells (infection detected in less than 1% of the animals), despite viral infection could be confirmed by RT-qPCR (see also Supplementary Table S4). Contrary, almost half the nematodes showed fluorescence in a control sample of plate-infected JU1580 validating the sensitivity of the assay (Ashe et al., 2013; Frézal et al., 2019) (Table S8). Furthermore, we used FISH on plate-infected IPR reporter strains (*pals-5::*GFP and F26F2.1::GFP in an N2 background) and observed that most nematodes that showed OrV FISH signal also showed IPR gene expression (Figure S6). About 43% of the individual nematodes in the population showed IPR expression (for both reporter strains). But many individuals that showed IPR gene expression did not show OrV FISH signal (Figure S6). More specifically, 21% of the animals that showed *pals-5* expression did not show OrV stained cells (Table S8). For the F26F2.1 reporter strain even 95% of the animals showed F26F2.1 expression but no OrV stained cells (Table S8).

**Supplementary Text S2.** ***IPR expression under heat stress***

IPR genes become differentially expressed upon prolonged heat stress providing thermotolerance in N2 (Reddy et al., 2017, 2019; Panek et al., 2020). Although the response of N2 and CB4856 to different temperatures has been well investigated, the role of IPR genes in heat stress has not been specifically compared between N2 and CB4856. We investigated the transcriptional IPR response to OrV heat stress in N2 and CB4856 by re-analyzing previously described data with a focus on the *pals*-genes (Jovic et al., 2017, 2019). The transcriptomes of heat shocked and recovering nematodes were compared to the control treatment. We found that IPR genes became differentially expressed after heat-shock and during recovery in both N2 and CB4856 (Supplementary Figure S7). The results demonstrate that the IPR is overall more active in CB4856 than in N2 in a completely unrelated experiment and show that the IPR can be activated in CB4856 upon heat-stress.

**Supplementary Table S1**. ***Genes involved during OrV infection*** *– Overview of C. elegans genes involved in OrV infection described in literature. Data is obtained from* (Félix et al., 2011; Ashe et al., 2013; Sarkies et al., 2013; Bakowski et al., 2014; Sterken et al., 2014; Tanguy et al., 2017; Chen et al., 2017; Jiang et al., 2017; Reddy et al., 2017, 2019; Le Pen et al., 2018; Sowa et al., 2019; Frézal et al., 2019; Sandoval et al., 2019).

*Table is provided as an .xlsx file*

**Supplementary Table S2.** ***Population genetic properties per pals-gene*** – *Conservation (genes are considered conserved when the vast majority of CeNDR strains contains < 25 polymorphisms), haplotype number and Tajima's D value per pals*-*gene. Genes that are upregulated upon infection in* (Chen et al., 2017) *are indicated.*

*Table is provided as an .xlsx file*

**Supplementary Table S3.** ***Haplotypes of pals-22 and pals-25 per CeNDR strain*** *– Haplotypes were manually inspected based for sequence coverage of pals-22 and pals-25 in the CeNDR database. The haplotype groups were noted for both genes per strain and divided into five categories: 'N2-like', 'CB4856-like', 'WN2002-like', 'CX11315-like' or 'ECA701-like'.*

*Table is provided as an .xlsx file*

**Supplementary Table S4.** ***Fluorescent signals observed in infected nematodes*** *– Summary of the phenotypes that were observed after OrV FISH staining in the strains N2, CB4856, ERT54, ERT71, and JU1580. The total number of inspected nematodes is shown and the number of nematodes showing either a green fluorescent signal (IPR reporter gene + GFP), red fluorescent signal (OrV RNA1.1) or both signals.*

*Table is provided as an .xlsx file*

**Supplementary Table S5**. ***Microarray probe binding to the CB4856 genome*** *– A) BLAST alignment of C. elegans CB4856 sequence (PJRNA275000) to microarray probes (C. elegans (V2) Gene Expression Microarray 4X44K slides) that detect the pals-genes. Probes are considered to align correctly when the there is more than 95% overlap on the correct chromosome. B) N2 and CB4856 DNA hybridization to the microarray probes genes that are differentially expressed based on genotype, treatment or their interaction. A ratio of 0 indicates there is absolutely no difference between N2 and CB4856 hybridization. The full dataset was previously published in* (Volkers et al., 2013) *and can be found online (ArrayExpress E-MTAB-8126). C) BLAST alignment of C. elegans CB4856 sequence (PJRNA275000) to microarray probes (C. elegans (V2) Gene Expression Microarray 4X44K slides) detect the IPR genes. Probes are considered to align correctly when the there is more than 95% overlap on the correct chromosome. D) BLAST alignment of C. elegans CB4856 sequence (PJRNA275000) to microarray probes (C. elegans (V2) Gene Expression Microarray 4X44K slides) that detect the differentially expressed genes from the linear model investigating the terms genotype, treatment and genotype x treatment. Probes are considered to align correctly when the there is more than 95% overlap on the correct chromosome. E) N2 and CB4856 DNA hybridization to the microarray probes genes that are differentially expressed based on genotype, treatment or their interaction. A ratio of 0 indicates there is absolutely no difference between N2 and CB4856 hybridization. The full dataset was previously published in* (Volkers et al., 2013) *and can be found online (ArrayExpress E-MTAB-8126).*

*Table is provided as an .xlsx file*

**Supplementary Table S6**. ***Output linear models*** *– A) The linear correlation between genotype, treatment, the interaction between genotype and treatment and gene expression for the N2 and CB4856 30 hours post (mock) infection. B) The linear correlation between viral load and gene expression within infected samples per genotype for the N2 and CB4856 30 hours post (mock) infection. C) The linear correlation between development, genotype, treatment and gene expression for N2, JU1580 and CB4856 infected samples that were isolated 1.5-30 hours post infection.*

*Table is provided as an .xlsx file*

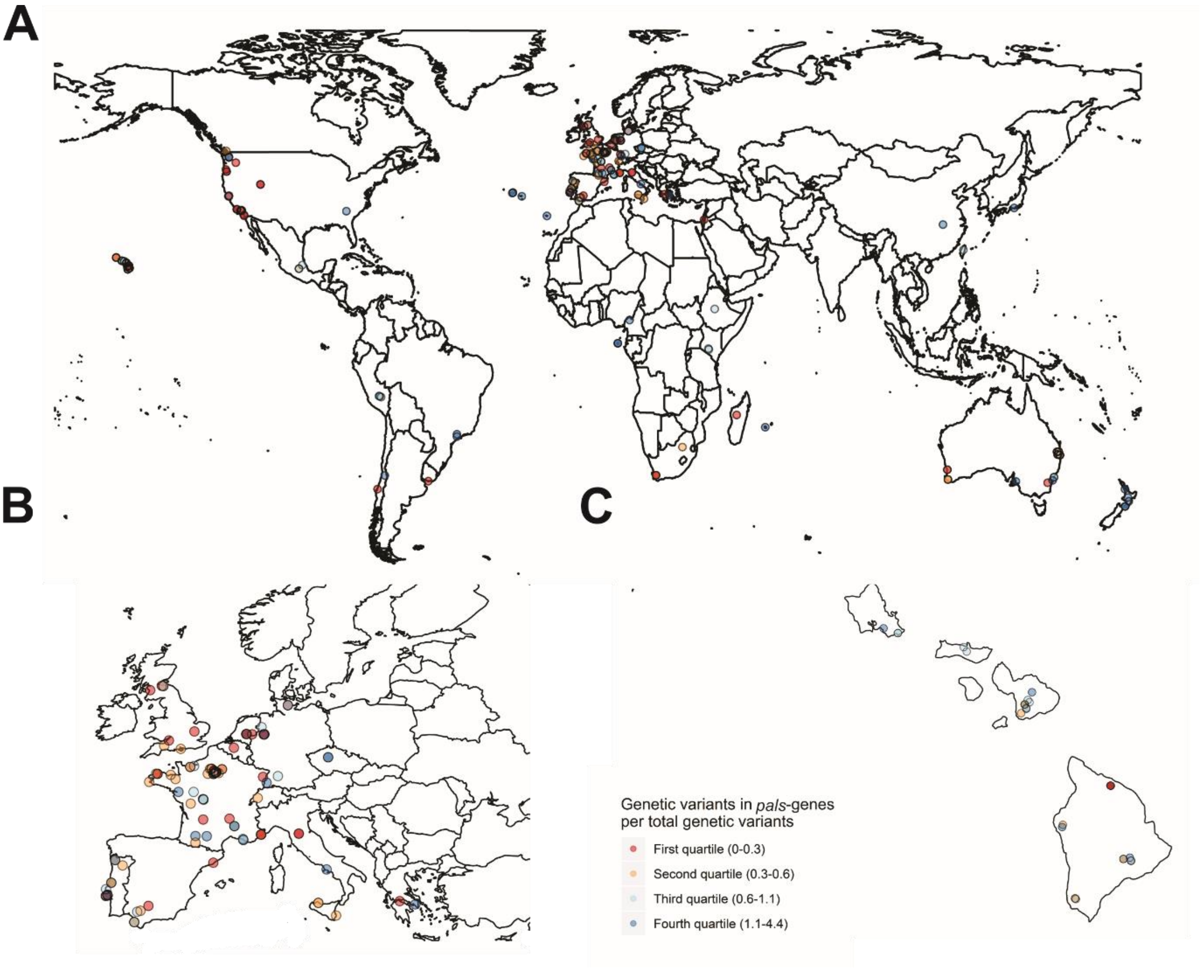

**Supplementary Figure S1.** ***Geographical distribution of natural variation within the C. elegans pals-gene family*** – *A) Location of CeNDR strains worldwide. The amount of natural variation (%) within the pals-pathway is indicated by the color of the dot. All natural strains have been grouped in a quantile (the first quantile exhibits least natural variation in the pals-pathway, the fourth exhibits most natural variation). B) Zoomed in representation of Figure S1A of the strains collected in Europe. C) Zoomed in representation of Figure S1A of the strains collected on Hawaii.*

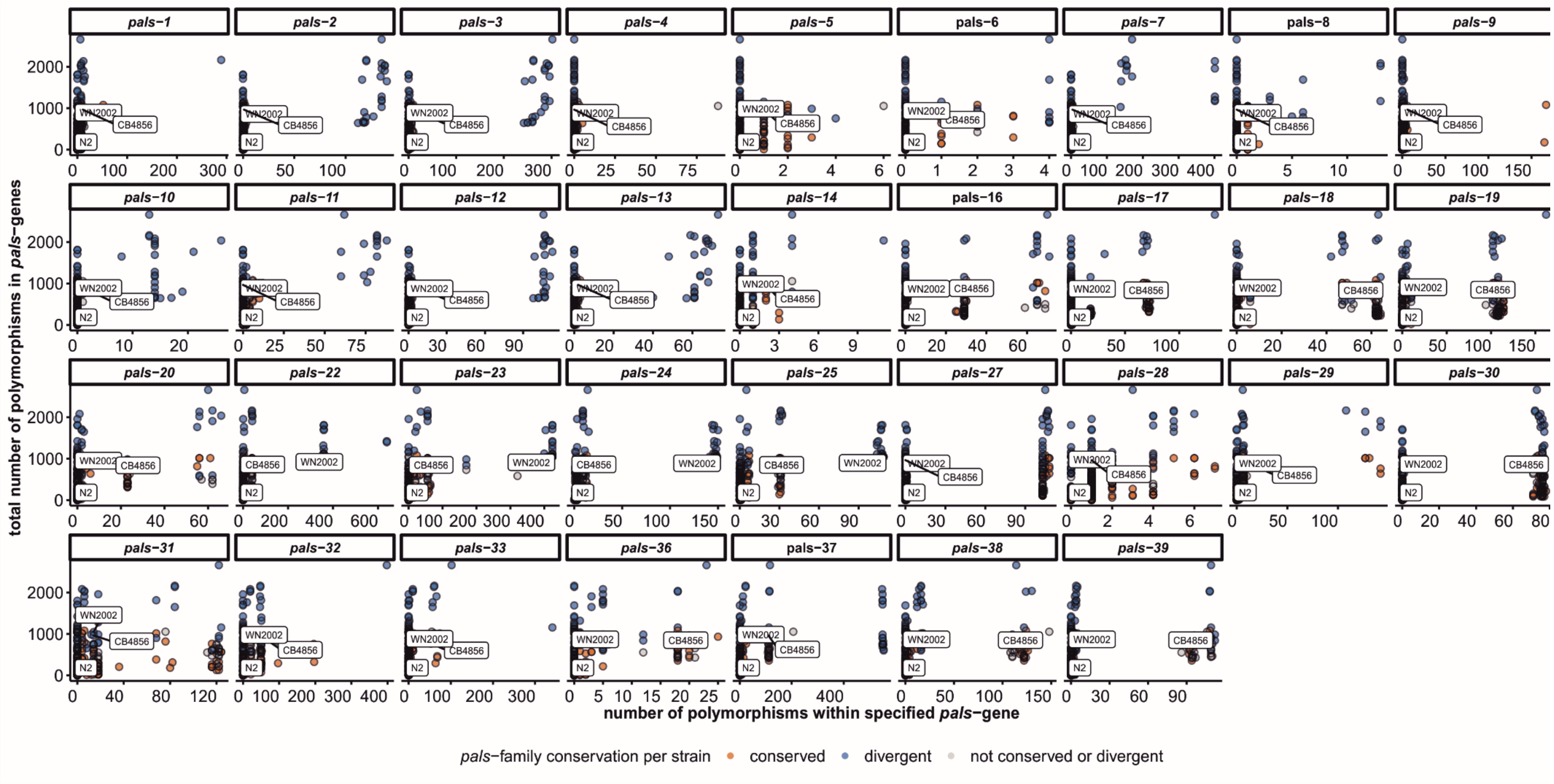

**Supplementary Figure S2.** ***Genetic variation per pals-gene per C. elegans strain*** *– The total number of SNPs within the pals-gene family is plotted against the number of known SNPs per pals-gene. Each dot represents a strain of the CeNDR database and the color of the strain indicates if the pals-family within this strain is depleted or enriched in polymorphisms as determined by the chi-square test (FDR < 0.0001).*

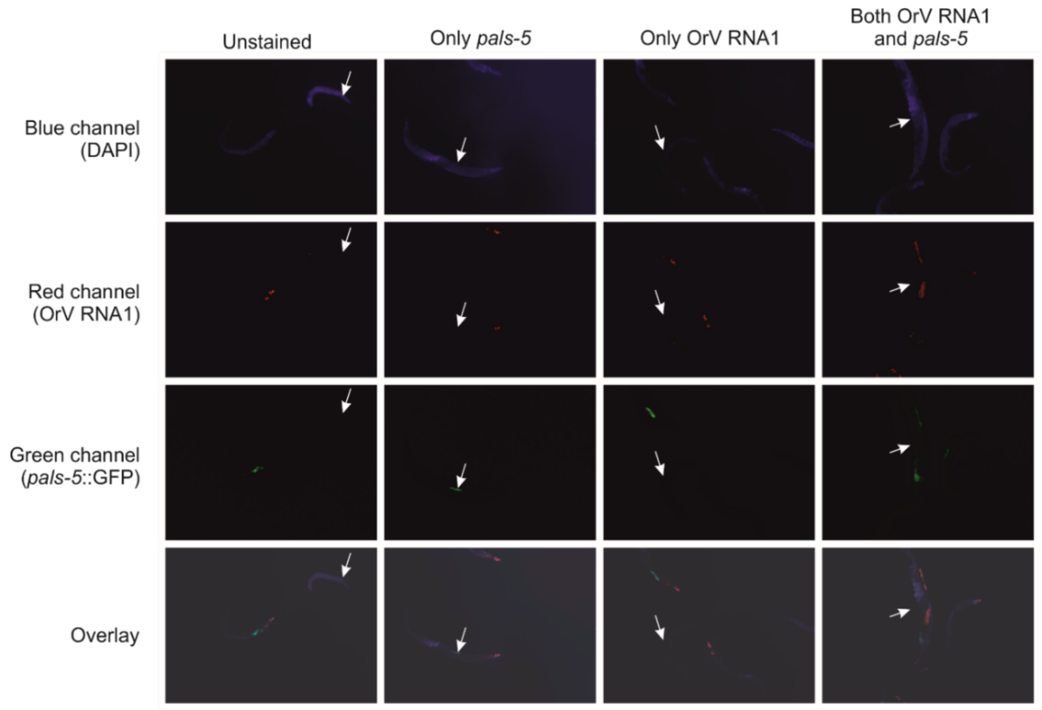

**Supplementary Figure S3.** ***Fluorescent in situ hybridization (FISH) of ERT54*** *– Examples of different phenotypes observed after OrV FISH staining in the ERT54 reporter (pals-5::GFP) strain. After staining plate-infected ERT54 nematodes, individuals expressing pals-5 and containing OrV infected cells were observed. These stainings could show in the same individual but were also observed separately. Different fluorescent channels show DAPI nuclear staining (blue), (intestinal) OrV RNA1 signal as stained by FISH (red), expression of the reporter pals-5::GFP (green). The exposure time in the red channel was fixed to 1s, for other channels the exposure times were set automatically. All nematodes show red fluorescence due to the myo-2::mCherry reporter in the pharynx.*

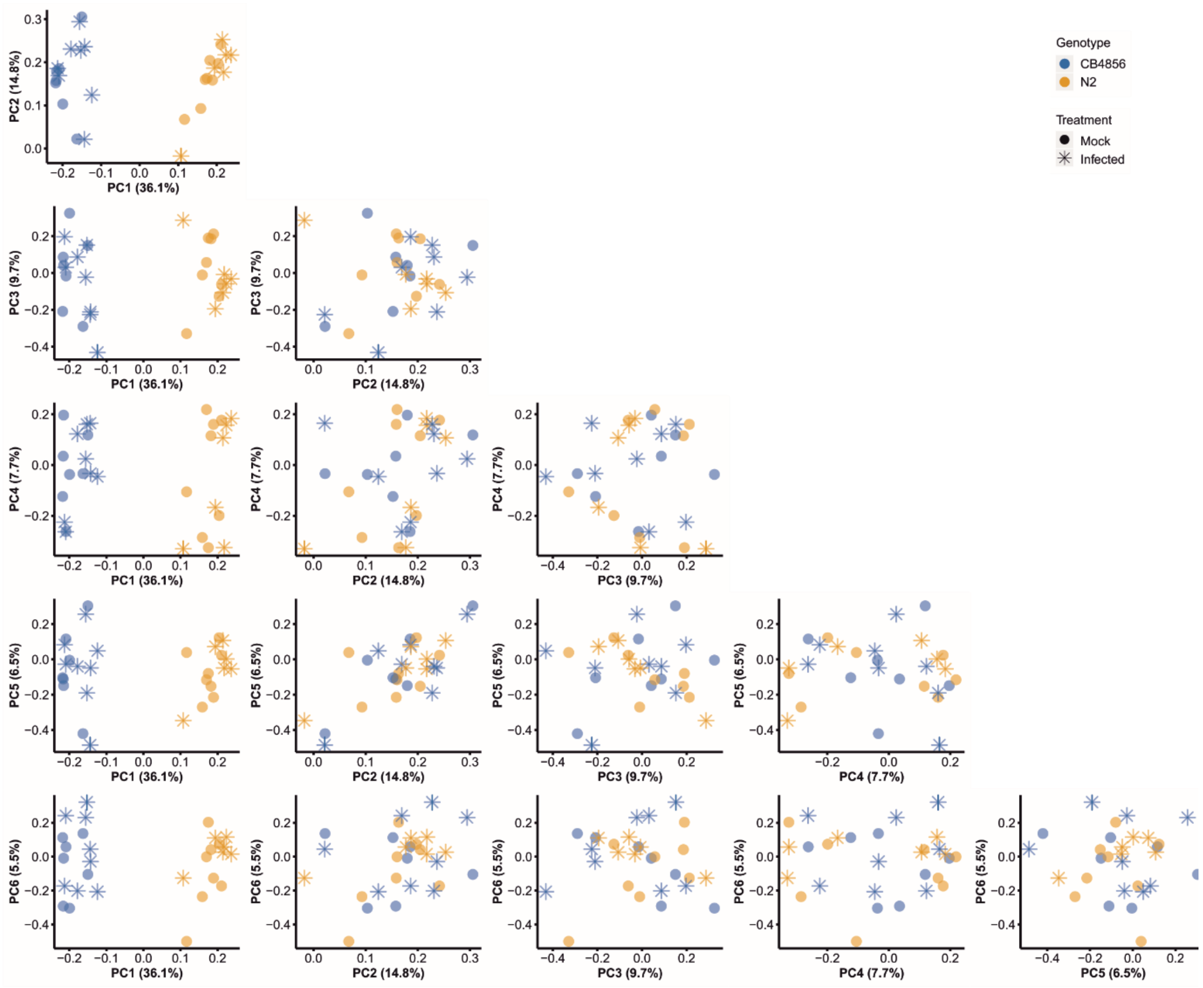

**Supplementary Figure S4**. ***Principal component analysis for gene expression in (un)infected C. elegans N2 and CB4856*** *– Principal component analysis for the gene expression data obtained for the nematodes that were infected 26 hours post bleaching and collected 30 hours post infection. The six PC axes that explain at least 5% of the total variation are shown, numbered from 1 (explaining 36.1% of variance) to 6 (explaining 5.5% of variance). Each dot resembles the location of a sample on these axes. The genotype (N2 or CB4856) is indicated by color and the treatment (mock or OrV infection) is indicated by shape.*

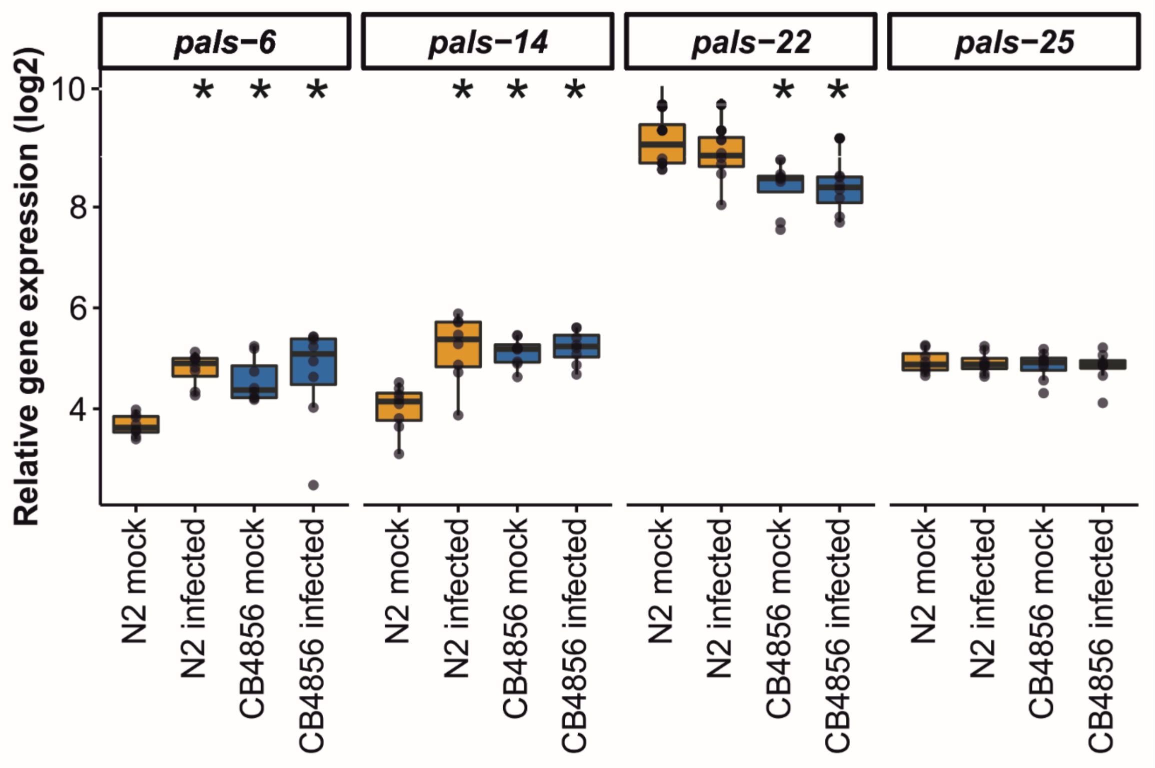

**Supplementary Figure S5**. ***Gene expression of pals-6, pals-14, pals-22, and pals-25 in the 30 hours exposure assay determined via RT-qPCR*** – *Box-plots of relative gene expression patterns for pals-6, pals-14, pals-22, and pals-25 which were determined using RT-qPCR. Measurements were performed on the same 32 samples that were used in the microarray analysis of (mock) infected N2 and CB4856. Each dot represents the expression within a sample. The expression of N2 mock was compared to N2 virus, CB4856 mock and CB4856 virus measurement per gene using a student t-test (*p < 0.05).*

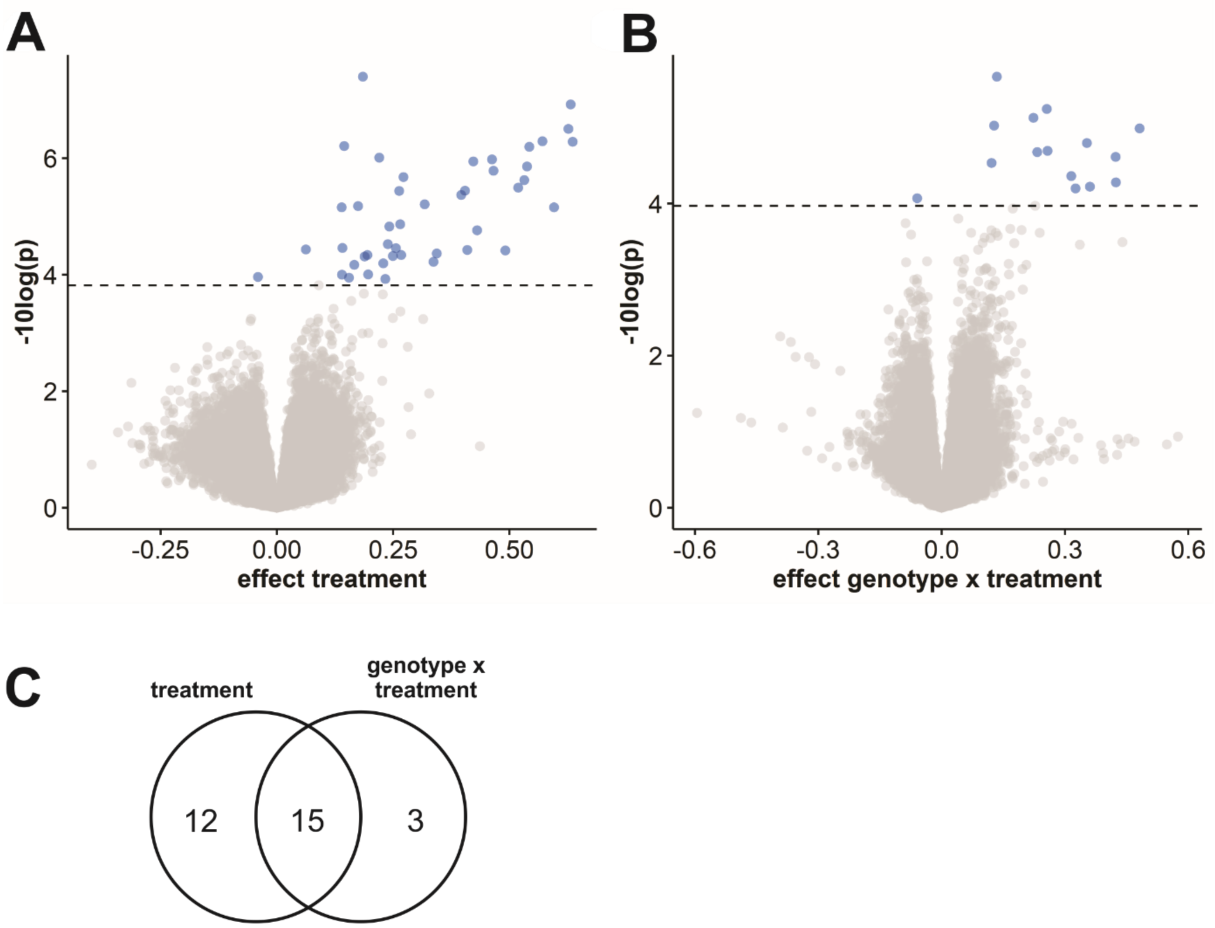

**Supplementary Figure S6**. *Differentially expressed genes upon OrV infection – A) Volcano plot showing the effect of treatment (OrV infection) on global gene expression patterns. Microarray spots (blue) above the FDR threshold (dotted line) were considered differentially expressed. B) Volcano plot showing the effect of the interaction between treatment (OrV infection) and genotype (N2 or CB4856) on global gene expression patterns. Microarray spots (blue) above the FDR threshold (dotted line) were considered differentially expressed. C) Venn diagram indicating the number of genes that are differentially expressed and associated with an effect treatment (OrV infection) and/or genotype x treatment.*

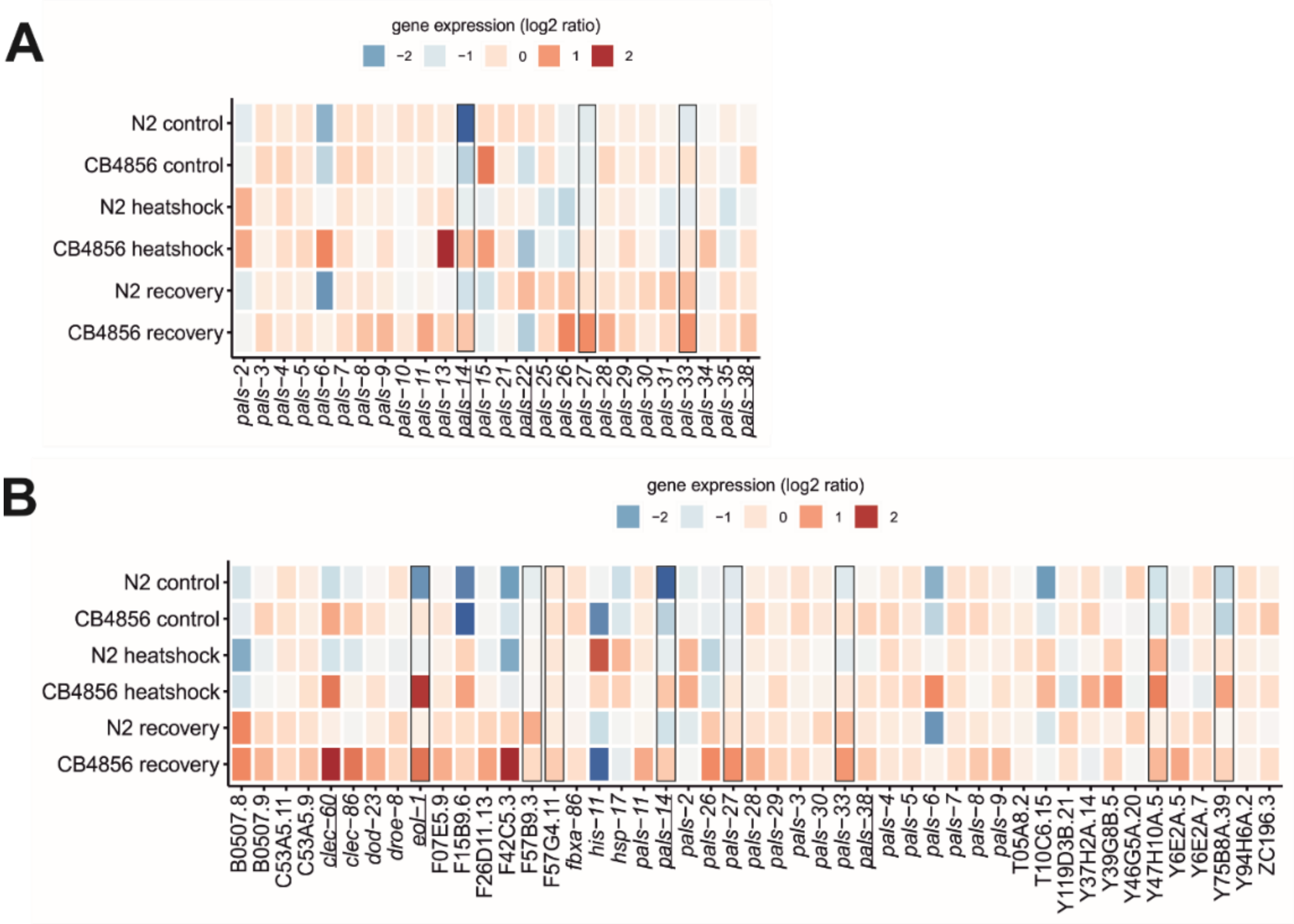
A
gene expression (log2 ratio)
−2 −1 0 1 2
N2 control
CB4856 control
N2 heatshock
CB4856 heatshock
N2 recovery
CB4856 recovery
pals−2 pals−3 pals−4 pals−5 pals−6 pals−7 pals−8 pals−9 pals−10 pals−11 pals−13 pals−14 pals−15 pals−21 pals−22 pals−25 pals−26 pals−27 pals−28 pals−29 pals−30 pals−31 pals−33 pals−34 pals−35 pals−38
B
gene expression (log2 ratio)
−2 −1 0 1 2
N2 control
CB4856 control
N2 heatshock
CB4856 heatshock
N2 recovery
CB4856 recovery
B0507.8 B0507.9 C53A5.11 C53A5.9 clec−60 clec−86 dod−23 droe−8 eol−1 F07E5.9 F15B9.6 F26D11.13 F42C5.3 F57B9.3 F57G4.11 fbxa−86 his−11 hsp−17 pals−11 pals−14 pals−2 pals−26 pals−27 pals−28 pals−29 pals−3 pals−30 pals−33 pals−38 pals−4 pals−5 pals−6 pals−7 pals−8 pals−9 T05A8.2 T10C6.15 Y119D3B.21 Y37H2A.14 Y39G8B.5 Y46G5A.20 Y47H10A.5 Y6E2A.5 Y6E2A.7 Y75B8A.39 Y94H6A.2 ZC196.3

***Supplementary Figure S7****. Gene expression of pals-genes and IPR genes before and after heat-shock – A) Heat-map showing the log2 intensities of pals-genes in N2 control, N2 heat-shock, N2 recovery, CB4856 control, CB4856 heat-shock and CB4856 recovery conditions B) Heatmap showing the expression of IPR genes in log2 intensities in N2 control, N2 heat-shock, N2 recovery, CB4856 control, CB4856 heat-shock and CB4856 recovery conditions. This dataset is described by (Snoek et al., 2017; Jovic et al., 2019). Underlined genes showed significant (basal) expression differences based on genotype (FDR < 0.05), whereas squares indicated the genes where treatment had a significant effect (FDR < 0.1). Log2 ratios are based on the average expression of the gene of interest in the overall dataset. Therefore, the log2 ratios per experimental group indicate the deviation from the average value. A subset of the pals-genes, namely the pals-genes that are also IPR genes (defined in (Reddy et al., 2019)) is depicted twice to facilitate direct comparison to non-IPR pals-genes and non-pals IPR genes.*

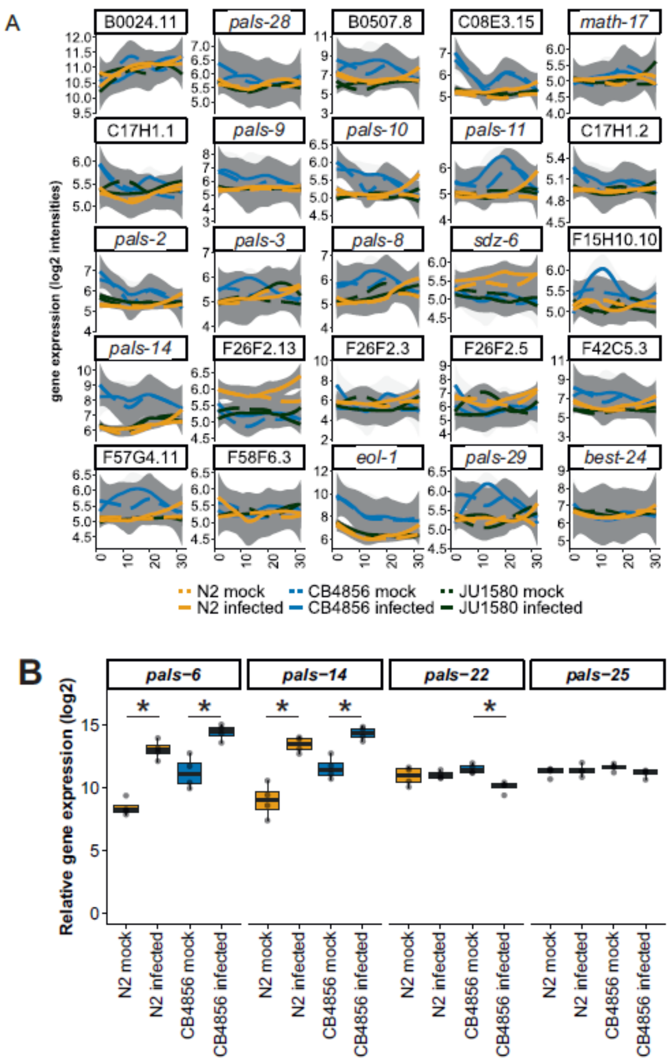

**Supplementary Figure S8.** ***Analysis of C. elegans expression patterns for OrV-response genes over time and long-term pathogenic exposure*** *– A) Gene expression patterns for N2 mock, N2 infected, CB4856 mock, CB4856 infected, JU1580 mock, and JU1580 infected nematodes of the 30 genes responding to OrV infection that were found in dataset of infected N2 and CB4856 nematodes (infected at 26 and collected at 56 hours post bleaching). The lines represent the fit through the data (by loess) and the grey area around a curve represents the 95% confidence interval, genotypes are indicated by colors and mock infection is indicated by dashed lines. B) Box-plots of relative gene expression patterns for pals-6, pals-14, pals-22, and pals-25 measured with RT-qPCR after exposure to OrV. Measurements were performed on the same 4 N2 and CB4856 samples that were obtained by continuous exposure to either mock-conditions or 50µL OrV. Each dot represents the expression within a sample. The expression of N2 mock was compared to N2 virus and the expression of CB4856 mock was compared to the CB4856 virus measurement per gene using a student t-test (*p < 0.05).*

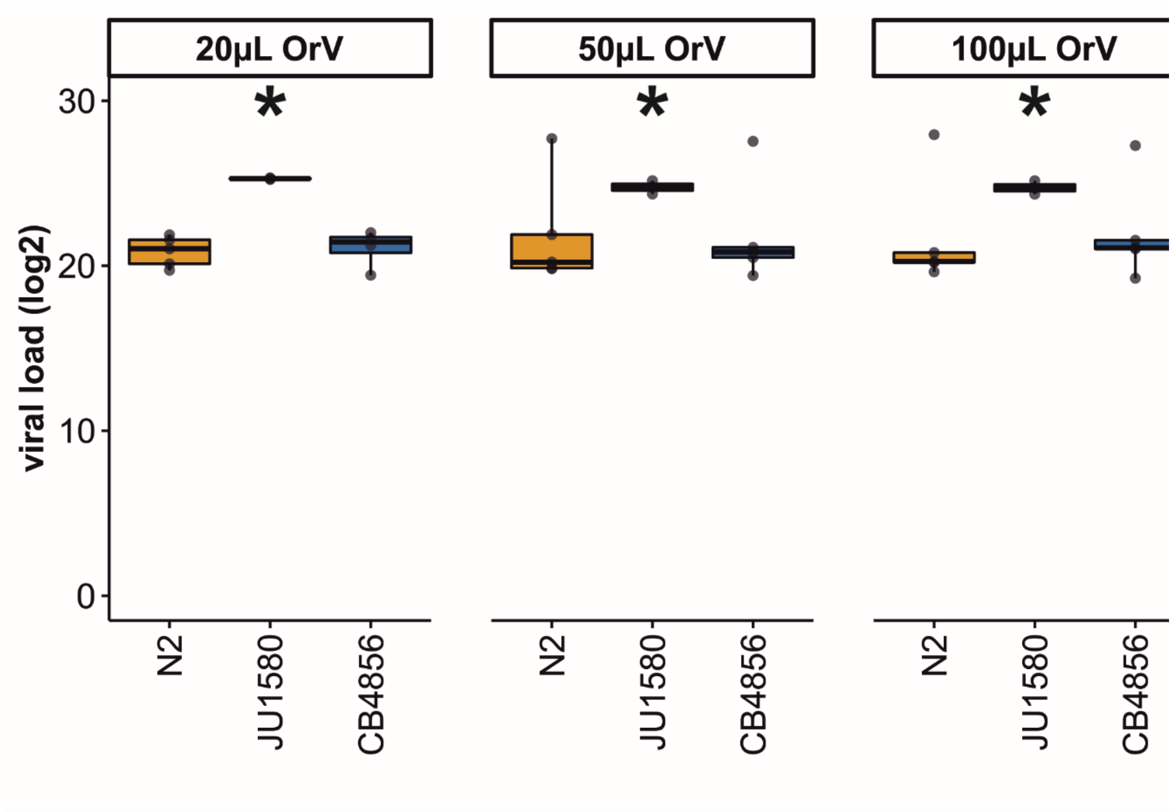

**Supplementary Figure S9**. ***Viral loads of C. elegans N2, CB4856, and JU1580 after 4 days of ex***
*or 100µL OrV to an NGM plate with three young adults followed by 4 days of incubation (4 bio*
*measuring viral loads ($log_2$) by RT-qPCR (student t-test; * $p < 0.05$ compared to N2). Boxplots show*
*bar indicating the median.*
